# Oxidative stress induces *E. coli* aryl polyene expression, sensitizing the bacterial stress response and modulating the redox environment of innate immune cells

**DOI:** 10.64898/2026.01.21.700819

**Authors:** Rachel L. Markley, Isabel Johnston, Viharika Bobba, Srishti Ithychanda, Genevieve Mann, Emma Dester, Kelsey Ouyang, Brianna P. Matthew, Jan Claesen

**Author notes:** Corresponding author: Jan Claesen.

## Abstract

Aryl polyenes (APEs) are specialized polyunsaturated outer membrane lipids that protect their producers from oxidative stress and contribute to biofilm formation. APEs are produced by an abundant biosynthetic gene cluster (BGC) family conserved across Gram-negative bacterial clades. The APE biosynthesis pathway involves 11 different enzymes and cumulates in the attachment of APEs to an anchor molecule in the Gram-negative outer membrane. Unlike most other small molecule BGCs, the APE BGC does not contain a dedicated regulatory gene that controls production of its metabolically costly compounds. Building from our prior observations of APEs role in acute oxidative stress protection, we here use a uropathogenic *Escherichia coli* (UPEC) strain to show that APE expression conveys a potential competitive advantage characterized by increased early-stage growth, sensitization of the bacterial oxidative stress response, and dampening of the redox stress of innate immune cells after *in vitro* infection. Our data indicate that APEs could act as a UPEC fitness factor, and in future work we aim to study their contribution to overall bacterial pathogenicity and survival, as well as how APEs could facilitate the transition from an oxygen poor environment such as the gut to the oxygen rich environment of the urinary tract.

**Importance:** Bacterial pathogens use various mechanisms to achieve a competitive advantage under harsh conditions such as upon interaction with their host. We studied the function of aryl polyenes (APEs), specialized polyunsaturated fatty acids in the outer membrane, in the context of a uropathogenic *E. coli* strain. APE expression is induced by an oxidative environment and contributes to early-stage growth and sensitization of the oxidative stress response. Furthermore, APE-expressing *E. coli* dampens the intracellular oxidative milieu of target host phagocytes. These findings suggest a role for APEs as a fitness factor, and create opportunities to study their *in vivo* function, and explore them as a potential drug target.

## Introduction

Reactive Oxygen Species (ROS) and Reactive Nitrogen Species (RNS) –collectively referred to as RNOS– are important molecular signaling molecules. They are vital to the survival of both bacteria and their host [1–4]. The host will generate RNOS through several NAPDH oxidases, and secondary peroxidases including neutrophil myeloperoxidase will further modify these RNOS molecules to signals that function to prevent infection by pathogens and opportunistic pathobionts [5]. In bacteria, RNOS signaling is involved in metabolic switching, such as during transition from anaerobic respiration to aerobic respiration. This process is initiated by catalase-peroxidase KatG, followed by specialized oxidases including cytochrome bd-II, and overall resulting in a metabolic advantage [6]. In addition to their role in autocrine and paracrine signaling in the host at low concentrations, RNOS serve a second role as potent antimicrobial agents when produced in large quantities by the innate immune system [7]. Hydrogen peroxide (H_2_O_2_) is a particular ROS species that can be found in the micromolar range in human urine, and is highly elevated in the blood and urine of patients with sepsis [8–11]. Because of their bactericidal activity, high levels of RNOS result in selective pressure in the microbiome, and provide opportunities for bacteria with enhanced fitness to outcompete other community members, leading to dysbiosis and associated pathologies [6, 12–14].

To cope with dramatic redox changes, bacteria have evolved several regulatory and quenching mechanisms that can act acutely as well as over an extended period [15–17]. Moreover, bacteria living in biofilms have enhanced survival chances because of their encapsulation in matrix and expression of protective metabolites on their cell envelope [18, 19]. One such protective metabolite is encoded by the aryl polyene (APE) biosynthetic gene cluster (BGC) [19–21]. The APE BGC family is widespread among Gram-negative bacteria, including most pathogenic *Escherichia coli* such as uropathogenic *E. coli* (UPEC) strain CFT073 [21]. APEs are yellow pigmented carboxylic acids comprised of an aryl head group and a conjugated double bond system, which are acylated to a larger lipid in the bacterial outer membrane, and they are structurally reminiscent of pigmented virulence factors found in Gram-positives [19, 22]. For example, the golden membrane-bound carotenoid virulence factor staphyloxanthin protects *Staphylococcus aureus* from oxidative stress and cationic peptides and helps protect the bacteria from neutrophil-mediated killing [23, 24]. Interestingly, staphyloxanthin can provide cross-species protection in a community setting during a polymicrobial infectious process [25]. Like staphyloxanthin, APE expression confers resistance to reactive oxygen species and enhances biofilm formation in *E. coli* [19]. These APE-associated characteristics are important attributes of UPEC bacteria that must withstand rapid changes to osmolarity, pH and oxygen levels during their commensal and pathogenic lifestyles. APE BGCs do not typically contain a dedicated regulatory gene [21] and it currently remains to be determined what regulatory cascades control APE gene expression, or how bacterial redox defenses interact in concert with the presence of APEs. Yet, since APE BGCs are present in several model pathogens and commensals, their involvement in microbe-host interactions can be inferred from mining the literature. Transcriptome analysis of uropathogenic *E. coli* identified genes in the APE BGC as induced fitness factors during human urinary tract infection [26]. Likewise, genes from enterohemorrhagic *E. coli* O157:H7 are upregulated in the presence of host epithelial cells [27] and its ApeH biosynthetic enzyme is expressed upon human infection, and implicated as a potential virulence factor [28].

Here, we report that APE expression enhances early growth *in vitro* using recombinant *E. coli* TOP10, a K-12 derived strain that lacks an endogenous APE BGC. We show that this growth advantage recapitulates for a constitutively expressing UPEC CFT073 APE^+^ strain. The early growth advantage in the presence of RNOS appears independent of the type of oxidative agent, and it is likely linked to oxidation of the APE metabolites, thereby protecting their producers. Oxidation of APE^+^ cells sensitizes expression of *oxyS*, allowing *E. coli* CFT073 to maintain redox homeostasis, which in turn could facilitate enhanced early growth. Finally, we investigate the potential implications for this RNOS mediation using an *in vitro* host-pathogen system, and show that innate immune phagocytic cells dampen their oxidative response when infected with APE-expressing bacteria. Taken together, our results suggest that APEs could act as a fitness factor for Gram-negative bacteria that harbor the BGC.

## Results

### APE expression confers an early growth advantage in *E. coli* exposed to oxidative stress

We previously showed that APE increases survival in a lab strain *E. coli* containing the cloned BGC from the uropathogenic strain CFT073 upon acute exposure to millimolar concentrations of H_2_O_2_ [19]. Here, we investigate the effects of APE expression under conditions of chronic exposure to RNOS stress generated by hydrogen peroxide (H_2_O_2_) and 3-Morpholinosydnonimine (SIN-1)(Fig. 1). Growth of *E. coli* Top10 heterologously expressing the APE BGC from a cosmid vector (TOP10^APE+^) is compared to an empty vector control strain (TOP10^EV^), which is naturally devoid of an endogenous APE BGC. Chronic exposure to 1 mM H_2_O_2_ results in a growth defect for TOP10^EV^ compared to the vehicle control (Fig. 1A), characterized by a two-fold increase in doubling time (Fig. 1B). In the TOP10^APE+^ strain, H_2_O_2_ exposure does not result in a similar growth defect (Fig. 1A) or doubling time (Fig. 1B), suggesting a protective role by APEs from an early growth stage. In contrast, no significant TOP10^APE+^ growth advantage was observed upon exposure to the superoxide and nitric oxide generating agent SIN-1 (Fig. 1C). In line with this observation, the doubling time during early growth did not show significant difference between strains under SIN-1 induced RNOS stress (Fig. 1D). Taken together, these growth assays indicate that APEs confer an early growth advantage in a chronically oxidative H_2_O_2_ environment.

**Fig. 1.**
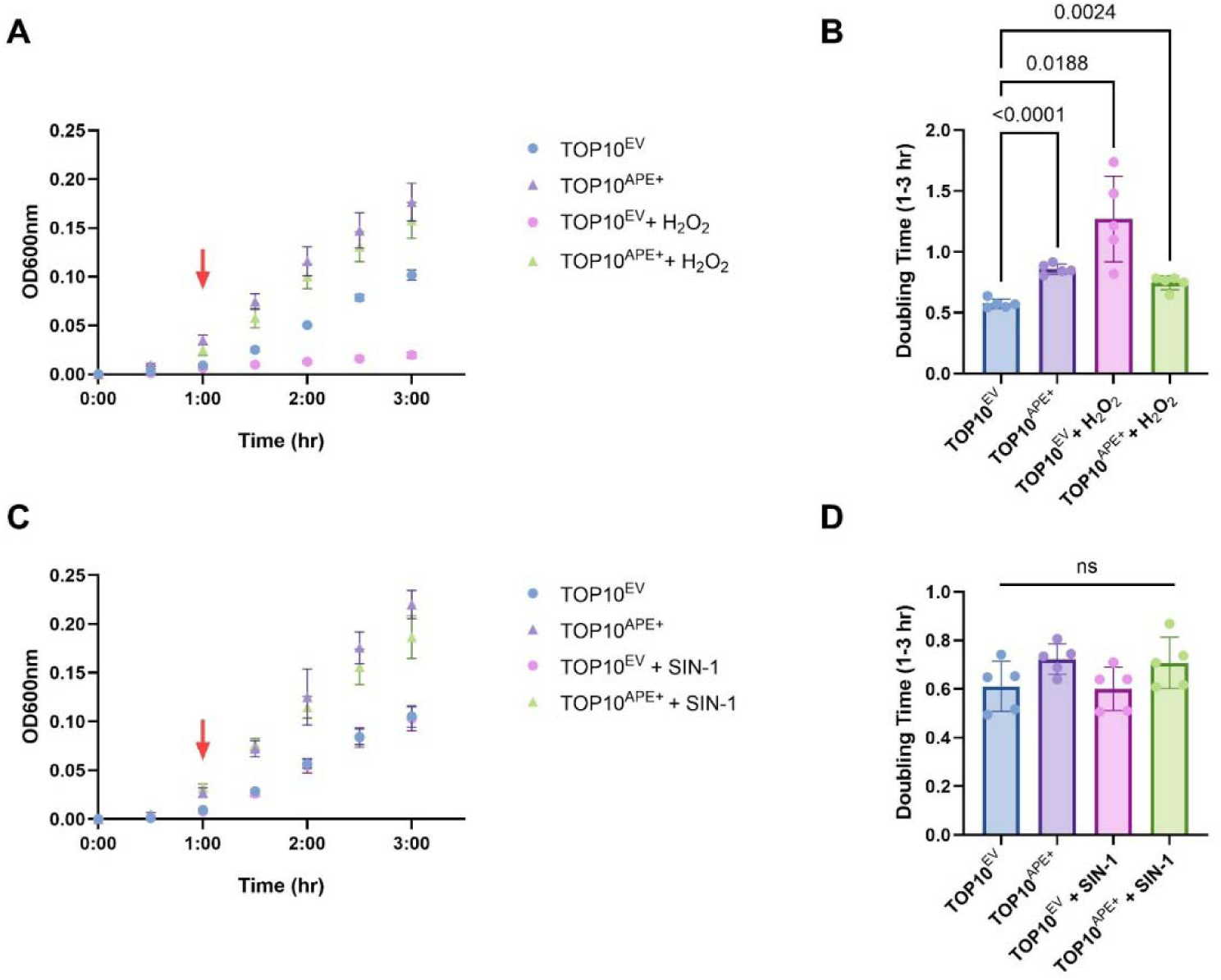
Growth dynamics of APE-producing *E. coli* Top10 exposed to chronic RNOS challenges. *E. coli* TOP10^EV^ and TOP10^APE+^ were cultured in a 96-well microplate at 37°C and growth was monitored by OD_600_ measurements. **(A)** Exposure to 1 mM hydrogen peroxide (H_2_O_2_) causes a growth defect for TOP10^EV^, but not TOP10^APE+^. **(B)** Quantification of bacterial growth using doubling time upon H_2_O_2_ exposure (T = 1-3 hr). **(C)** Growth curves for *E. coli* TOP10^EV^ and TOP10^APE+^ exposed to various concentrations of 3-Morpholinosydnonimine (SIN-1). **(D)** Quantification of bacterial growth using doubling time upon SIN-1exposure (T = 1-3 hr). *P-*values shown were calculated using a Lognormal Brown-Forsythe and Welch ANOVA test with a Dunnett’s T3 multiple comparison test. (n=5 biological replicates per group)

### RNOS exposure reduces APE pigmentation

We observed that the prolonged exposure to oxidative agents causes a discoloration of the yellow pigmented APE^+^ bacteria over time, likely due to oxidative destruction of the conjugated double bond system. To measure oxidative breakdown of APEs, we performed a bulk lipid extraction of the cell envelopes of TOP10^APE+^ bacteria. Given that this lipid extraction procedure yielded highly concentrated samples, we were necessitated to use supraphysiological levels of RNOS in the following bleaching experiments. We exposed the cell extract to various RNOS and monitored absorbance at 441 nm to assess changes to the APE pool [21]. There was a decrease in yellow pigment upon addition of 3 M H_2_O_2_ and the extract was significantly bleached after 24 hours, as compared to the vehicle control (Fig. 2A-C). Similarly, exposure to SIN-1 resulted in decreased pigmentation after 24 hours (Fig. 2D-E).

**Fig. 2.**
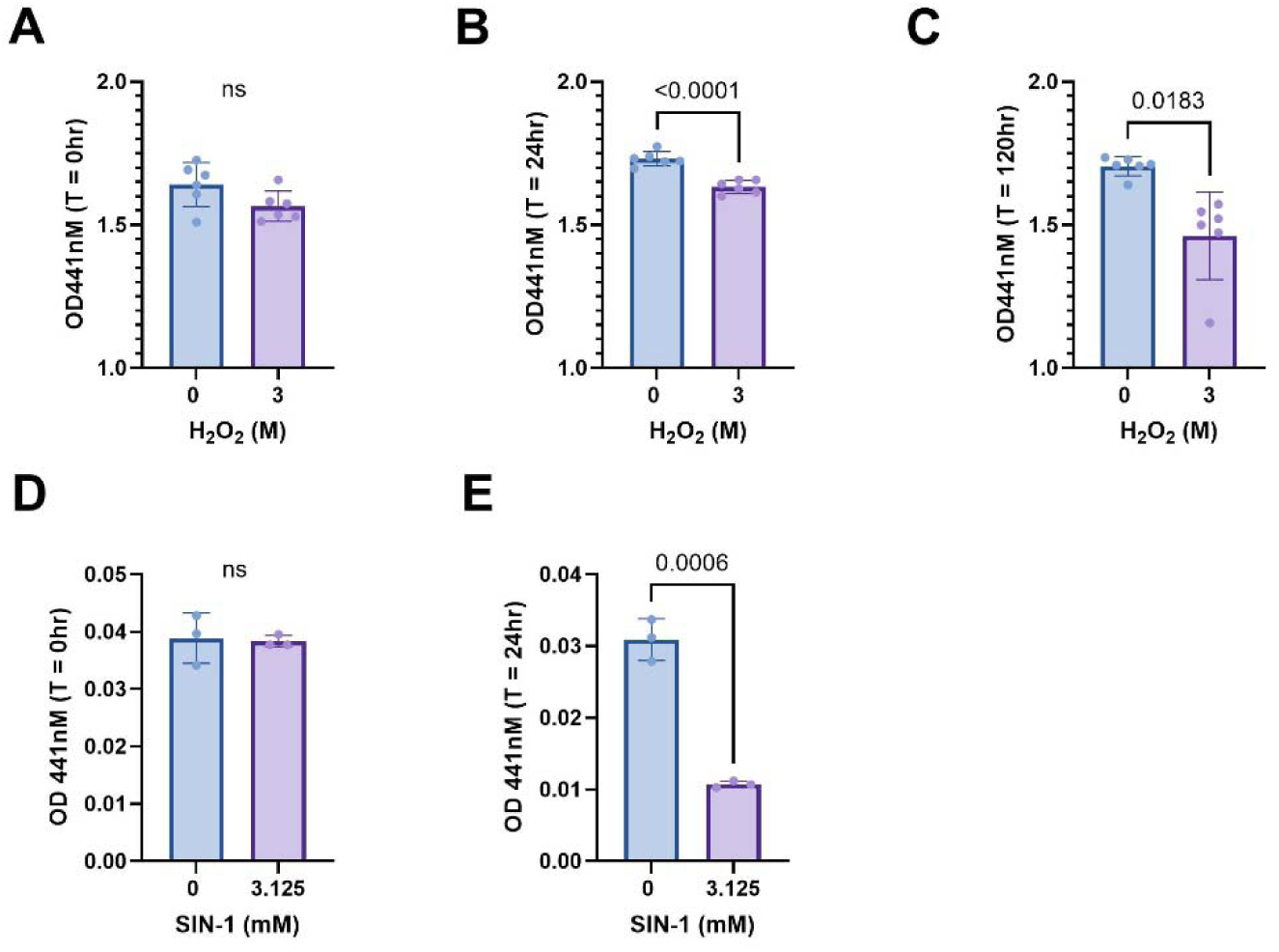
Bleaching of APE^+^ cell extracts exposed to oxidative agents. **(A)** Absorbance values at OD 441 nM upon H_2_O_2_ exposure at T = 0hr, **(B)** T = 24 hr, and **(C)** T = 120 hr (5 days). **(D)** Oxidation of Top10^APE+^ extracts by SIN-1 at T = 0hr, and **(E)** T = 24hr. *P-*values shown were calculated by an unpaired Lognormal Welch’s t-test. (n=3-6 biological replicates per group)

### Redox pressure induces APE gene expression and sensitizes the oxidative stress response in *E. coli* CFT073

We next examined the effects of an oxidative environment on APE expression in *E. coli* CFT073, a UPEC strain and native APE BGC host. Congruent with our observations with *E. coli* Top10, a CFT073^APE+^ ectopically expressing strain exposed to H_2_O_2_ had a selective growth advantage with a shorter lag phase, compared to the CFT073^EV^ control (Fig. 3A) as measured by optical density at time points before 8 hr. We determined bacterial viability after 5 hours of growth and as expected found a reduction in CFT073^EV^ CFUs in cultures exposed to H_2_O_2_. By contrast, we observed an increased survival in the CFT073^APE+^ strain, which even doubled at 0.75 mM H_2_O_2_ as compared to the vehicle control (Fig 3B). The early-stage growth advantage conferred by CFT073^APE+^ was confirmed by the increased total viability of APE-expressing bacteria (CFU/ml) exposed to H_2_O_2_ compared with vehicle-treated controls within the CFT073^APE+^ strain (Fig. 3C).

**Fig. 3.**
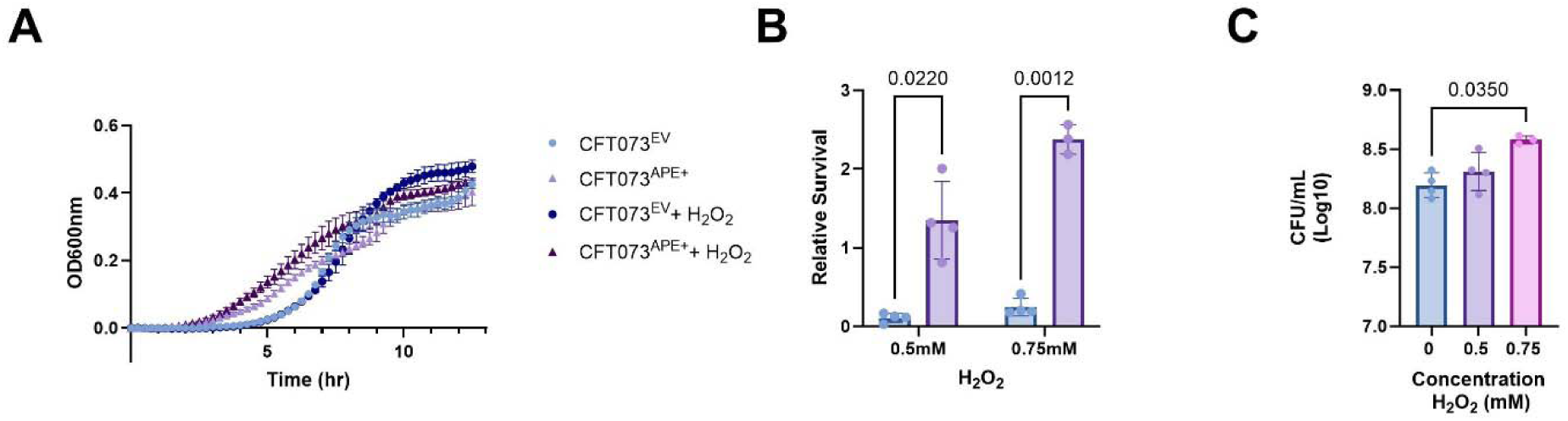
APE expression confers an early-stage growth advantage in *E. coli* CFT073 exposed to H_2_O_2_. **(A)** *E. coli* CFT073^EV^ and CFT073^APE+^ were cultured in a 96-well microplate at 37°C and growth was monitored by OD_600_ measurements. **(B)** Relative survival was calculated by dividing bacterial counts for each condition by the average count recovered at 0 mM H_2_O_2_. *P-*values were calculated by Mixed-effects model with the Geisser-Greenhouse correction, followed by a Š□dák’s multiple comparisons test, with individual variances computed for each comparison. (C) Total viability of CFT073^APE+^ measured as CFU/ml at 5 hr after inoculation. *P-*values were calculated by Kruskal-Wallis ANOVA, followed by a Dunn’s multiple comparisons test. (n=3-4 biological replicates per group)

To assess whether the increased protection was related to APE production, we measured expression of select genes in the APE BGC of *E. coli* CFT073^EV^ in response to H_2_O_2_ challenge. We observed modest H_2_O_2_-dependent relative mRNA level increases of the early biosynthesis gene *apeO* (encoding 3-ketoacyl-ACP-synthase) and a similar trend for *apeR* (encoding a different 3-ketoacyl-ACP synthase), as well as an increase in mRNA levels for the attachment gene *apeD* (encoding lysophospholipid acyltransferase). We did not observe any expression differences, for example for *apeB*, which encodes the methyltransferase involved late in APE carboxylic acid biosynthesis (Fig. 4A-E) [19]. Repeating this experiment using SIN-1 as a stressor, we corroborated increases in *apeO* and *apeD*, and observed a non-significant increasing trend in *apeR* expression. We reasoned that the modest increases in APE BGC expression might not fully explain the observed fitness advantage and extended our analysis to general antioxidant mechanisms. When assaying transcriptional responses at a more acute timepoint, 15 minutes after exposure with 1mM of H_2_O_2_, we did not note increased APE BGC expression. To our surprise, mRNA levels of *oxyR*, a sensor and positive regulator of the *E. coli* H_2_O_2_-inducible regulon [29], were induced specifically in the CFT073^APE+^ strain, but not CFT073^EV^ (Fig. 5A). This differential induction was amplified in the expression levels of the downstream small untranslated regulatory RNA *oxyS* (Fig. 5B). Similarly, the mRNA levels of the OxyR-dependent catalase-peroxidase *katG* increased specifically in CFT073^APE+^ after 15 minutes of H_2_O_2_ exposure (Fig. 5C). In summary, our results indicate that APE BGC expression is triggered by chronic oxidative stress, and that APEs sensitize their producers’ *oxyR*-dependent stress response.

**Fig. 4.**
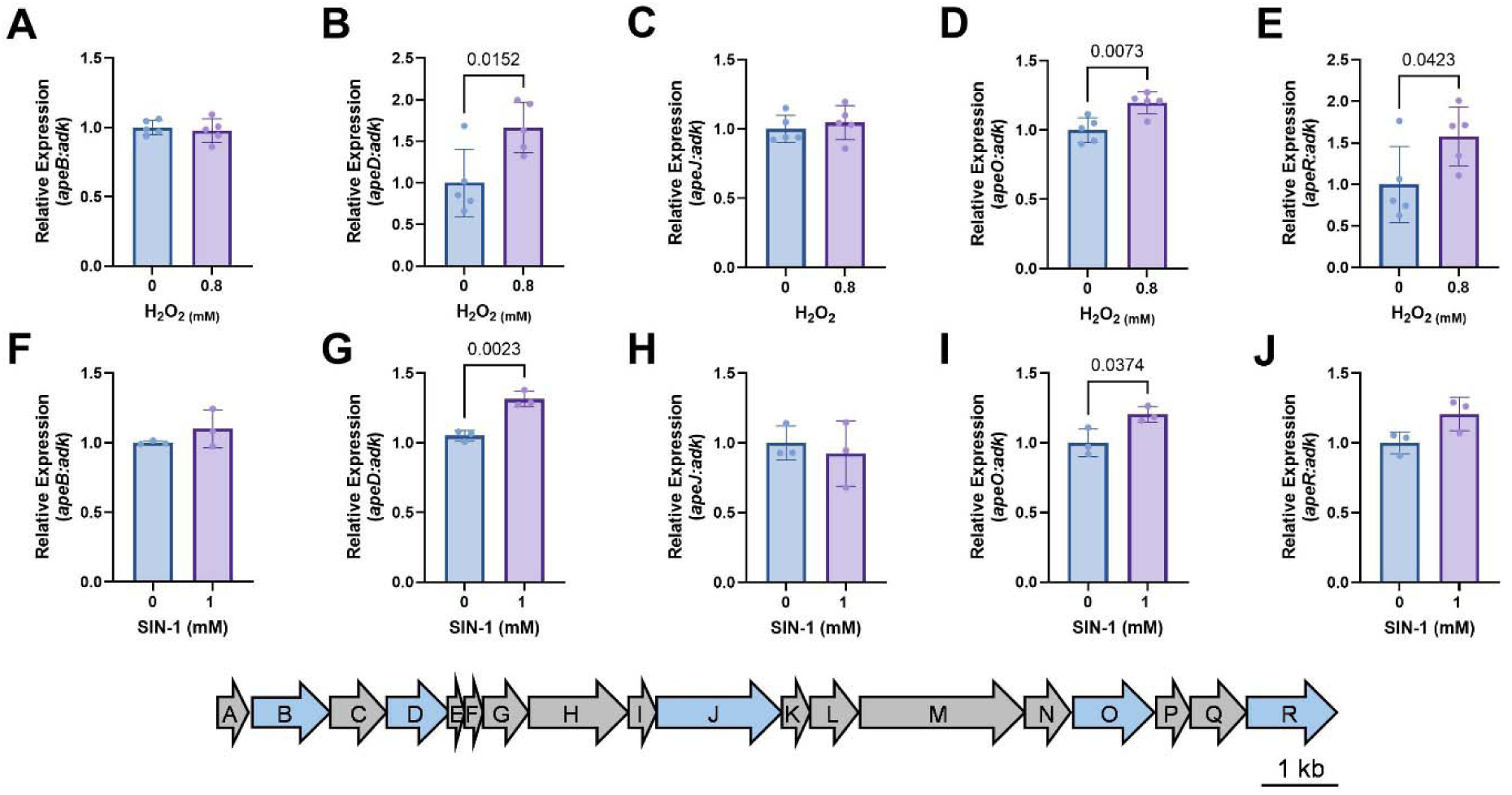
mRNA expression analysis for select genes in the *E. coli* CFT073^EV^ APE BGC after 24 hrs chronic exposure to H_2_O_2_ (A-E) or SIN-1 (F-J). (A,F) *apeB* (SAM-dependent methyltransferase), (B,G) *apeD* (lysophospholipid acyltransferase), (C,H) *apeJ* (glycosyltransferase/acyltransferase), (D,I) *apeO* (3-ketoacyl-ACP-synthase), (E,J) *apeR* (3-ketoacyl-ACP-synthase). Gene expression was normalized to *adk* and *p-*values were calculated by an unpaired Lognormal t test for all except (F) which included the Welch’s correction due to differences in the significant variance. The genes for which expression was analyzed are indicated in blue on an arrow representation of the APE BGC. (n=3-5 biological replicates per group)

**Fig. 5.**
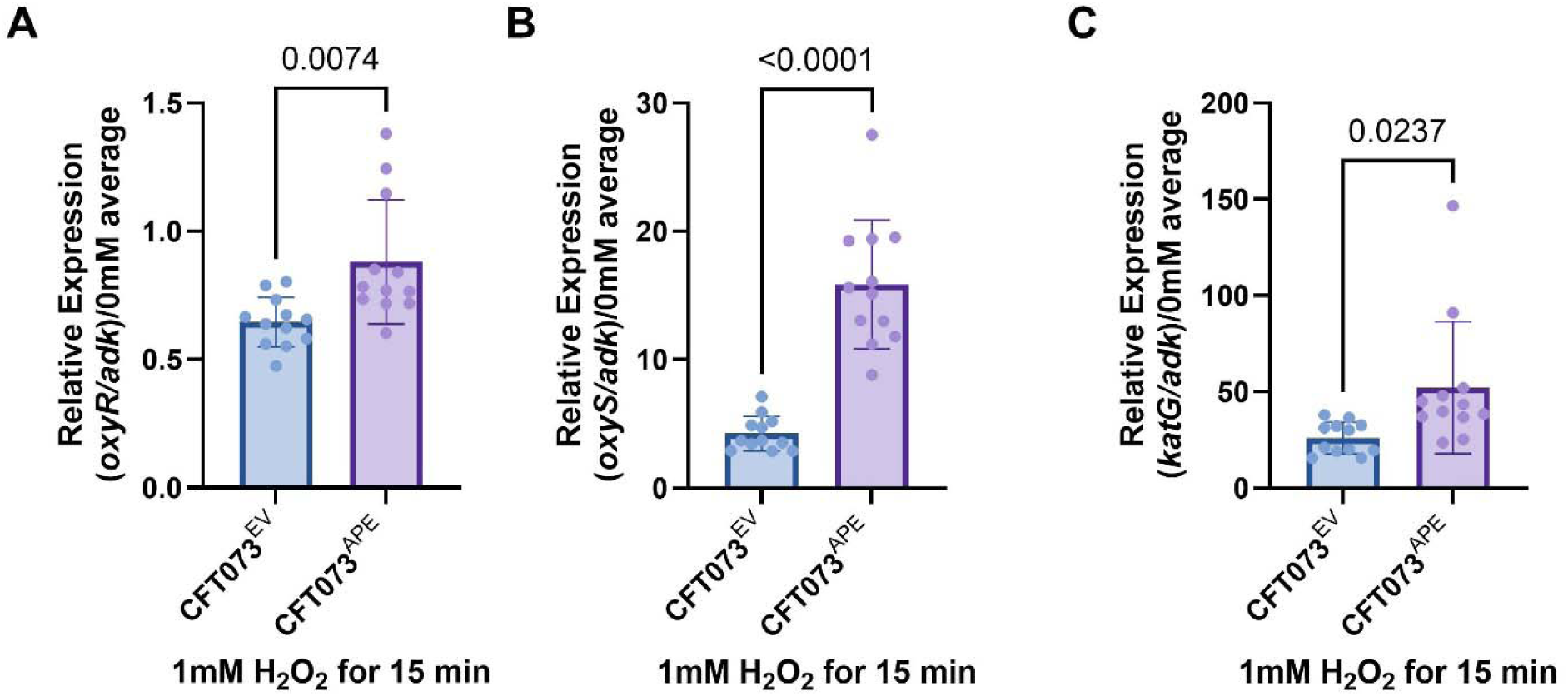
APE BGC expression sensitizes *E. coli* CFT073’s oxidative stress response. Relative fold change for (A) *oxyR*, (B) *oxyS*, (C) *katG* (n=12 biological replicates per group). *P-*values were calculated from an unpaired t-test with Welch’s correction.

### Phagocytes infected with *E. coli* CFT073^APE+^ have a dampened oxidative stress response

Host phagocytes contribute to the early stages of the innate immune response to infection by acutely producing oxidative agents, a process referred to as ‘respiratory burst’ [30, 31]. This process is in part driven by myeloperoxidase (MPO), an enzyme responsible for utilizing H_2_O_2_ with available anions to make antimicrobial molecules such as HOCl [32]. We investigated the impact of *E. coli* APE expression on the oxidative environment of murine macrophage-like RAW 264.7 cells using the CellROX assay for ROS quantitation. RAW 264.7 cells were infected with *E. coli* CFT073 EV, KO or APE^+^ at a multiplicity of infection (MOI) of 10:1 and their oxidative stress levels were compared to uninfected controls by flow cytometry analysis. Infection of RAW 264.7 cells with CFT073^APE+^ caused a ~4-fold reduction in cells positive for the CellROX stain as compared to control macrophages or compared to infection with CFT073^EV^(Fig. 6A). We corroborated these results by performing an analogous infection in primary human neutrophils and assaying for MPO expression. In line with our results with the RAW 264.7 cells, fewer neutrophils (identified by CD11b and CD16 expression) that expressed MPO were present in the samples infected with CFT073^APE+^ compared to CFT073^EV^ or CFT073^KO^ infection or uninfected controls (Fig. 6B). Moreover, the median fluorescent intensity of those neutrophils that did stain positive for MPO was lower than that of cells infected with CFT073^EV^ or CFT073^KO^ (Fig. 6C). These data suggest that bacterial APE expression can have an extrinsic effect on the phagocyte intracellular environment by reducing RNOS production. Hence, exposure to APE producing bacteria could diminish the population of immune cells which participate in RNOS-driven antimicrobial activities, which could have implications during an active infection.

**Fig. 6.**
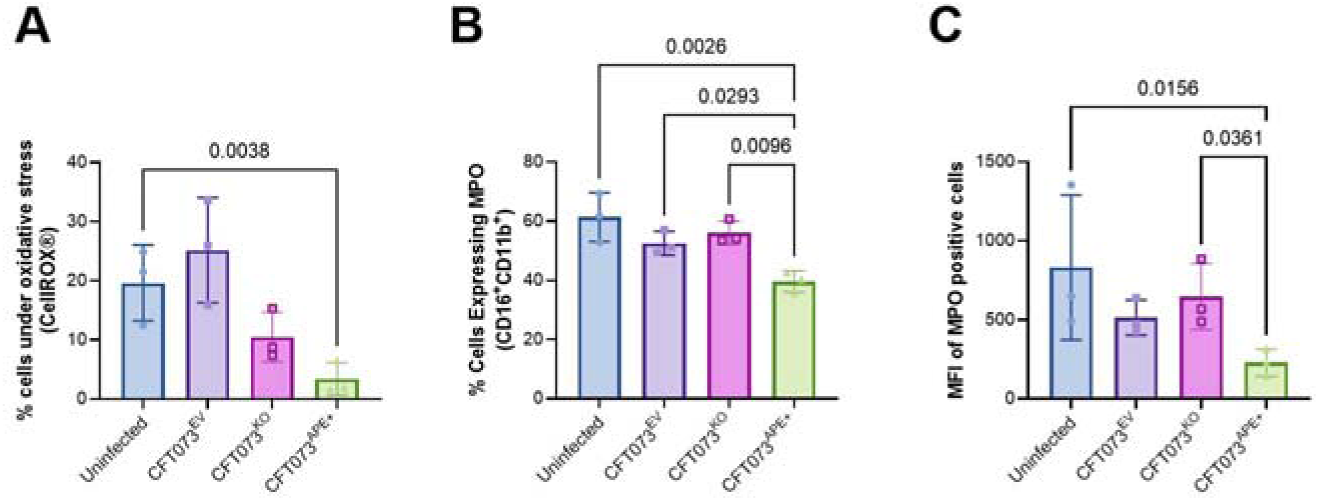
Oxidative stress response of host phagocytes infected with APE expressing bacteria. **(A)** Murine RAW 264.7 cells were infected with *E. coli* CFT073 EV, KO or APE^+^ and stained for ROS using the CellROX kit, followed by quantification by flow cytometry. **(B)** Freshly isolated human neutrophils were infected with *E. coli* CFT073 EV, KO or APE^+^ and stained for CD11b and CD16 to identify them as neutrophils and then intracellular staining for myeloperoxidase (MPO) was followed by quantification by flow cytometry. All statistics were lognormal ordinary ANOVA test with a Tukey’s multiple comparison test using a single pooled variance. (n=3 biological replicates per group)

## Discussion

The APE BGC is found widely distributed among Gram-negative bacterial species. We and others have previously implicated its role as a fitness factor, both by protecting from acute ROS challenge, as well as by increasing biofilm formation kinetics [19]. Here, we report a distinct early-log phase growth advantage of *E. coli* APE producers under selective pressure of RNOS, using recombinant strains of both the Top10 cloning strain as well as the UPEC CFT073 strain. One ramification of increased respiratory energy production during exponential growth is the increased generation of RNOS, which could lead to cellular damage. We postulate that APE expression could contribute to a fitness advantage that may offset the downstream damage of this increased oxidative pressure. It is of note that for both Top10 and CFT073, the APE^+^ strains have a baseline growth advantage compared to the EV strain in the absence of oxidative stress (Fig. 1A,C and Fig. 3A). Since the APE BGC is expressed from its native promoter(s), this could potentially be due to differences in plasmid copy number. Overall, this increased baseline growth advantage is counterintuitive, as the additional production of APEs would be expected to be metabolically burdensome for the cells, as reflected in the doubling times (Fig. 1B). Future research will focus on the dual nature of APE as its function might switch from a protective effect during early growth towards a more burdensome or even toxic role at later growth stages. It is possible that this change is coupled to metabolic switches in the cells, such as utilization of different amino acid substrates in the LB medium, the accumulation of acetate, rise of the pH or fluctuations in oxygen levels [33, 34]. Future work can dissect the relative contribution of these parameters, and we acknowledge that their relative *in vivo* importance might not become clear from growth studies in LB medium.

Oxidative stress can be harmful to a multitude of cellular molecules and structures, including proteins. A class of enzymes which are particularly sensitive to oxidative stress are iron-sulfur cluster containing proteins like formate dehydrogenase (Fdh) [35]. To mediate RNOS stress damage, *E. coli* has evolved various types of antioxidant systems. We found that constitutive expression of APE sensitizes the thiol-response element *oxyR* in *E. coli* CFT073. OxyR senses oxidated lipids in the cell envelope [36] and hence oxidation of APEs could potentially amplify this signal. We also observed increased expression of the downstream small regulatory RNA *oxyS,* as well as catalase-peroxidase *katG*. This robust antioxidant response in APE^+^ cells could in part explain the growth and survival advantages despite elevated levels of RNOS. In addition, an oxidative environment will increase APE gene expression, suggesting a diversion of bacterial metabolism to maintain APE expression. More studies will be needed to fully examine APE expression with different carbon sources to determine if RNOS stress from specific metabolic activities triggers APE expression.

Upon exposure to RNOS, the characteristic yellow APE pigmentation was bleached, suggesting the molecules may be acting as a sacrificial reactive oxygen sink to mitigate damage to other molecules and structures in their producers. This ability to diminish oxidative damage also has the potential to increase the survival of APE producing pathogenic bacteria in a host-associated context. In addition, we show an *in vitro* reduction in oxidative stress and MPO activity in murine RAW 264.7 macrophage-like cells and primary human neutrophils exposed to APE expressing *E. coli* CFT073. We therefore demonstrate that APE confers a multi-angled protective strategy against RNOS, providing direct protection, promoting more rapid microbial antioxidant expression, and exerting a dampening effect on RNOS produced by infected host cells. Taken together, this insinuates a potentially important survival advantage for APE producers in host-pathogen interactions and might explain why so many pathogenic Gram-negative bacteria have retained the APE BGC in their chromosome. In future studies, we will address the physiological implications of APE expression during host-pathogen interactions and determine if APE expression leads to increased pathology.

## Methods

### Bacterial strains and culture conditions

Bacterial cultures were maintained in liquid Luria-Bertani (LB, Novagen Millipore) growth medium (175 rpm, 37 °C), or on LB agar (BD Difco) at 37 °C, and supplemented with 50 µg/mL kanamycin and/or 100 μg/ml carbenicillin (Fisher BioReagents) when appropriate. *E. coli* TOP10 (Invitrogen, ThermoFisher Scientific), *E. coli* Top10 APE^+^ (containing pJC121), and *E. coli* CFT073 were described before. *E. coli* CFT073 APE^+^ was generated in this study by introducing pJC121 via electroporation [19, 37]. All bacteria used in the described experiments were cultivated in the presence of 50 µg/mL kanamycin to maintain their plasmids. Prior to use in all assays, bacteria were streaked fresh from a frozen glycerol stock and incubated on solid medium for 24 hrs. For growth assays, bacterial strains were normalized by suspension in LB broth to 10^7^ CFU before 1:1000 inoculation. All growth analysis were completed after collected OD_600_ values were first normalized to the starting OD_600_ value at T = 0hr. Growth assays were performed in a Biotek plate reader using the Gen 5 software (Synergy HT, Biotek/Agilent Systems), measuring absorbance at 600 nM.

### Crude APE^+^ cell envelope extraction and oxidation assays

Liquid cultures were grown at 37°C in LB broth for 5 days, pelleted by centrifugation (at 4000 × *g*), washed with phosphate buffered saline (PBS) and thrice with water before they were weighed and frozen. Equivalent masses of wet cell pellet were first extracted with acetone and then extracted twice in 2:1 acetone:MeOH (v/v) in a blender. Organic extracts were pooled and their volume reduced in a rotary evaporator prior to liquid-liquid extraction with an equal volume of diethyl ether. The organic phase was collected and evaporated to dryness for storage or reconstituted in MeOH for use in oxidation assays. Oxidations of the APE^+^ cell envelope extracts were carried out with hydrogen peroxide (Thermo Scientific), and 3-Morpholinosydnonimine (SIN-1; Cayman Chemicals) suspended in water (H_2_O_2_ and or MeOH (SIN-1). 441 nM absorbance measurements were carried out on a Biotek plate reader using the Gen 5 software.

### Gene expression analysis

Bacterial cells in (LB medium) were incubated 1:3 (v/v) in RNAprotect (Qiagen) for 5 min at RT. Next, cell pellets were obtained by centrifugation (10,000 × *g* for 10 min) and lysed using 10 mg/mL Lysozyme (Thermo scientific) in TE buffer for 10 min at RT. Total RNA was isolated using the RNeasy Plus kit (Qiagen) and 300–500 ng was used to synthesize cDNA with the qScript cDNA Synthesis Kit (Quanta bio) following the manufacture’s protocol. cDNA was diluted 1:5 to 1:10 and used as template in a quantitative PCR reaction with Fast SYBR Green Master Mix (Applied Biosystems). qPCR primers listed in Table 1 were designed based off the *E. coli* CFT073 complete genome sequence (GenBank: AE014075.1) [Welch 2002]. Expression data was collected and analyzed using the StepOnePlus Real-Time PCR System and StepOne Software v2.3 (Applied Biosystems). Relative quantification of gene expression was assessed using the 2^-ΔΔCT^ method using *adk* as a housekeeping gene.

**Table 1:**
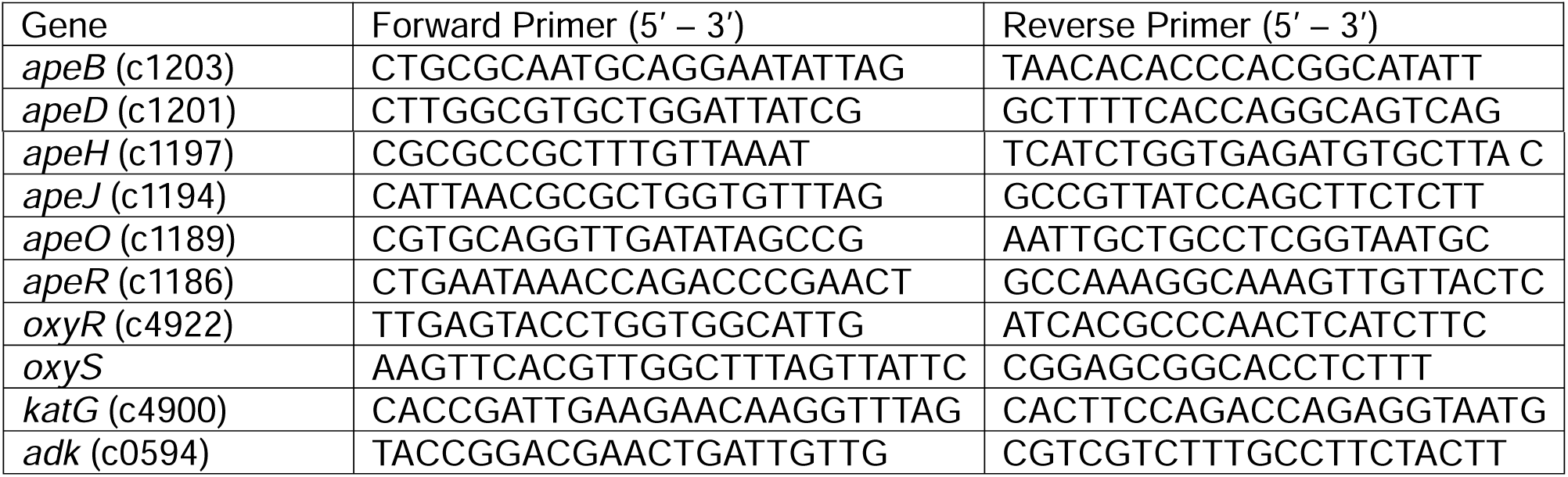
Primers used in this study.

### CellROX Assay

RAW 264.7 cells were plated in 6 well plates at 1 × 10^6 cells/well. The cells were maintained in DMEM High Glucose (HG) without phenol red with 5% Heat inactivated fetal bovine serum, 10mM HEPES, with 2mM L-glutamine in a 37°C, 5% CO_2_ and inoculated with rCFT073 diluted in DMEM without phenol red with kanamycin to 10^7 CFU under the aforementioned culture conditions. Cells were spun at 300 × *g* for 10 min and incubated at 37*C for 2 hours. The cells were then rinsed with PBS and trypsinized to transfer to an Eppendorf tube where they were washed with and resuspended in PBS. Cells were stained with Live/Dead Fixable Blue™ (Invitrogen) and resuspended in PBS with 1% bovine albumin. CellROX™ Green Flow Cytometry Assay Kit (Life Technologies) was completed using the manufacture’s protocol. Cells were fixed eBiosciece™ Intracellular Fixation Buffer (Invitrogen). Flow cytometry analysis was completed by the Flow cytometry core at CCF using the LSRII with the BD FacsDiva software version 8.0.1 set to acquire 5000 events per sample. Results were analyzed via FlowJo™ v10.8 Software (BD Life Sciences) the gating strategy used FSC-A X SSC-A to eliminate debris. FSC-H X FSC-A was used to eliminate doublets and UV450-A negative cells provided a population of live cells that were then further gated on stained cells (Blue 515-A) to determine the % population under ROS stress. Unstained and single stained controls were used to aid in gating strategy.

### Myeloid Peroxidase Assay

Neutrophil isolation from the buffy coat was achieved through density centrifugation as described previously [38]. Briefly the buffy coat was layered over LSM™ lymphocyte separation medium (MP Biomedials) and centrifuged at 500 × *g* for 35 min at 25°C. A neutrophil band was carefully removed and diluted in Hanks Buffered Saline Solution (HBSS) without calcium or magnesium. Cells were centrifuged at 350 × *g* for 10 min at 25°C. Red blood cells were lysed using the ACK Lysing Buffer (Gibco™) according to the manufacturer specifications. After lysis cells were washed with HBSS. Cells were infected in HBSS at multiplicity of infection of 10:1. Cells were washed with PBS and stained with Live/Dead Fixable Blue™ (Invitrogen) and resuspended in PBS with 1% bovine albumin. Neutrophils were blocked with Human TruStain FcX™ (Biolegend) and stained with anti-CD16-APC (3G8, Biolegend), anti-CD11b-PerCP-Cy5.5 (LM2, BioLegend), anti-CD45-Alex Fluor 488 (2D1, BioLegend). Cells were washed before fixation and permeabilization with eBiosciece™ Intracellular Fixation and Permeabilization Buffer (Invitrogen). Intracellular staining was carried out for anti-MPO-eFluor 450 (MPO455-BE6, Invitrogen). Flow cytometry analysis was completed by the Flow cytometry core at CCF using the LSRII with the BD FacsDiva software version 8.0.1 set to acquire 5000 events per sample. Results were analyzed FlowJo™ v10.8 Software (BD Life Sciences). Single cell controls and fluorescence-minus-one controls were used to aid in gating. Gating strategy was: 1.) FSC-H X FSC-A to eliminate doublets, 2.) FSC-A X SSC-A to eliminate debris, 3.) gating on UV450-A negative cells provided a population of live cells, 4.) CD45 positive cells were gated (Blue 515-A) with a low SSC-A. 5.) Cells were gated for double positive of CD11b/CD16 (Red-670 and Blue-710A). 6.) Gating cells positive for MPO (violet 450-A) a cell count and median fluorescence were calculated.

### Statistical Analysis

All graphs and statistical analyses were generated using the Graphpad Prism software, details on specific statistical tests applied are mentioned in the respective figure legends.

### Data availability statement

We did not generate any large datasets as part of this study; all data are available in the main text or can be obtained upon request.

## Acknowledgements

We thank Dr. Tom McIntyre and Dr. Rui Chen for providing buffy coat and for their intellectual support for neutrophil isolation. We also thank current and past members of the Claesen, Ahern and Hajjar labs and the Flow Cytometry Core at the Lerner Research Institute for their intellectual support and technical help. This work was supported by a National Institutes of Health (NIH) grant R01 AI153173 (J.C.), and in part by seed funding from the Cleveland Clinic Foundation (J.C.).

## Author contributions

R.L.M. and J.C. conceived and designed the experiments. R.L.M., I.J., V.B., S.I., E.D., K.O., G.M., B.M., and J.C. performed experiments or analyzed the subsequent data. R.L.M. and J.C. wrote the paper and handled visualization. All authors discussed the results and commented on the manuscript.

